# Flipped elevational pattern of pollination mode in tropical vs. temperate Americas

**DOI:** 10.1101/2022.03.04.483035

**Authors:** Agnes S. Dellinger, Ashley M. Hamilton, Carolyn A. Wessinger, Stacey Smith

## Abstract

**Aim:** Abiotic factors, such as temperature and precipitation, vary markedly along elevational gradients, and can in turn, shape key biotic interactions, such as herbivory and pollination. Despite the well-known effects of climatic conditions on pollinator activity and efficiency, we know little about the role of climate in pollinator shifts in animal-pollinated plants at broad geographic scales. Here we investigate patterns of altitudinal turnover in pollination mode across the Americas, with a focus on the most common pollinators (bees and hummingbirds). Specifically, we test Cruden’s classic hypothesis that plants are likely to shift to bird pollination at high elevations because endothermic pollinators are more reliable in cold and rainy conditions.

**Location:** Americas

**Time period:** Current

**Major taxa studied:** 2232 plant taxa from 26 clades

**Methods:** We collated information on pollination mode (1262 insect-pollinated, 970 vertebrate-pollinated) for the study taxa from the literature, and used GBIF occurrence data to estimate median distributions and bioclimatic attributes of each species. We used (phylogenetic) GLMMs to test for associations between pollination mode and ecogeographic variables.

**Results:** To our surprise, we found flipped elevational patterns of insect- and vertebrate-pollination strategies across latitudes, with vertebrate pollination dominating at high elevations in the tropics, but not in temperate zones. We term this pattern the ‘Tropical flip’. We recovered a strong association of vertebrate-pollinated plants with moist, forested habitats across latitudes, while insect-pollinated plants were often found in cool and dry or warm and moist conditions.

**Main conclusions:** Altitudinal gradients in temperature may not serve as a universal explanation for shifts among endothermic insect and ectothermic vertebrate pollination. Instead, strong abiotic niche differentiation among insect- and vertebrate-pollinated plants, along with competition for pollination niche space, has likely shaped the ‘tropical flip’.

## Introduction

Biotic interactions such as pollination are among the most prominent factors limiting, promoting and structuring organismal diversity on earth (Schemske et al. 2009). Studies of plant-pollinator interactions often focus on the functional relationships, such as the fit between flowers and pollinator mouthparts (Whittall & Hodges 2009) and the potential for functional trade-offs to drive pollinator shifts (Muchhala 2007). Nevertheless, pollinator shifts often have a geographic component due to spatial variation in pollinator abundance and efficiency (Dellinger et al. 2021). This pattern emerges at a macroecological scale as major groups of pollinating animals (e.g. hummingbirds, bees) vary in importance across latitudinal and altitudinal gradients (Schemske et al. 2009, Ashworth et al. 2015, Classen et al. 2015, McCabe & Cobb 2021, Dellinger et al. 2021). For example, bats, rodents and passerine birds are important pollinators particularly in tropical regions (Ratto et al. 2018), while insects and hummingbirds act as pollinators across latitudes. Similarly, diverse insect pollinator assemblages (i.e. bees, beetles, wasps, butterflies), common at low elevations, are gradually narrowed to bumblebee- and particularly fly-dominated pollinator communities at high elevations (Warren et al. 1988, Arroyo et al. 1982, Primack & Inouye 1993, Lefevbre et al. 2019, McCabe & Cobb 2021). Identifying the ecogeographic and climatic factors underpinning, and potentially driving these patterns, remains a major task.

Across the Americas, recurring evolutionary shifts between bee and hummingbird (but also bat or rodent) pollination have been documented in unrelated plant clades, occasionally resulting in large radiations of vertebrate-pollinated lineages in tropical mountains (Lagomarsino et al. 2016, Serrano-Serrano et al. 2018). In his groundbreaking work, Cruden (1972) hypothesized that such shifts between insects and vertebrate pollinators are driven by climate, providing a mechanistic explanation for the well-known geographic pattern. Specifically, his work demonstrated that the adverse abiotic climatic conditions prevalent in tropical montane environments (i.e., cool, moist, windy, cloudy) dramatically reduce the flower-visitation activity (and hence pollination efficiency) of bees, but not of hummingbirds (also see Dellinger et al. 2021). Clearly, bees, and many other small insects, are ectothermic, and their activity patterns are regulated by the ambient temperature, while vertebrates are endothermic and hence not restricted in their flower visitation activity to spells of sunny, warm weather (McCallum et al. 2013). Further, in contrast to insects, vertebrates may continue visiting flowers even in moderate rain, common in the changeable weather conditions of mountains (Lawson & Rands 2019). Temperature and precipitation hence have fundamentally different effects on the activity of insect and vertebrate pollinators (McCallum et al. 2013, Classen et al. 2015). These climate-induced reductions in bee-pollinator activity, along with reduced bee diversity and abundance (Perillo et al. 2021), likely cause the turnover from bee to vertebrate pollination moving up along elevational gradients (Thomson & Wilson 2008, Dellinger et al. 2021).

While Cruden’s altitude-driven pollinator shift scenario, formalized on data from tropical mountains, has formed a hallmark of pollination biology research for decades, some reports indicate that hummingbird pollination in temperate zones is unrelated to high-elevation ecosystems (Stebbins 1989). Instead, large bumblebees, which, similar to vertebrates, can regulate their own body temperature, act as important mid to high elevation pollinators (McCallum et al. 2013). At critically low temperatures (around 5°C mean annual temperature) at highest elevations, flies replace bumblebees as pollinators (McCabe & Cobb 2021). This pattern of elevational turnover in pollinator communities suggests that the simple model of altitude-driven insect-to-vertebrate pollinator shifts is not universally applicable. Instead, hummingbird-pollinated temperate plant species appear to be associated with slightly mesic, shady environments such as deep canyons (Grant & Grant 1968, Stebbins 1989). Such habitats may, just like tropical montane forests, represent environments of reduced bee pollinator efficiency, freeing pollination niche space for hummingbirds (Cruden 1972, “hummingbird habitats”, Stebbins 1989). Despite the fundamental importance of pollinator shifts in the diversification history of angiosperms, to our knowledge, no attempts have been made so far to systematically test Cruden’s altitude-driven bee-to-vertebrate pollinator shift scenario across the tropics and temperate zones, and identify the bioclimatic thresholds linked to such shifts.

Here, we address these research gaps by investigating ecogeographic patterns of shifts between insect (particularly bee, but also fly, moth, wasp, generalist) and vertebrate (bird, bat, rodent) pollination across 2232 plant taxa from 26 clades across the entire Americas. First, we test Cruden’s (1972) altitude-driven pollinator shift scenario to determine whether vertebrate pollination is consistently more common at higher elevations. Against expectations, we do not find support for this hypothesis: while vertebrate-pollinated species indeed occur significantly more often at high elevations than bee-pollinated species in the tropics, the pattern is flipped in temperate zones. Given these contrasting patterns with elevation, we next examined whether turnover from insect to vertebrate pollination is linked to specific climatic conditions that limit the activity of insects but not vertebrates (i.e. low temperature, high precipitation, high cloud cover). We find evidence of bioclimatic differentiation among pollination strategies, with vertebrate-pollinated species occurring at intermediate temperatures (10-20°C) under moist conditions in forested habitats, while insect-pollinated species are found in both warmer and wetter as well as cooler and drier conditions. Overall, our results are important in providing the hitherto broadest assessment of ecogeographic factors associated with shifts among insect and vertebrate pollination systems, and paint a new picture of interacting biotic and abiotic niche differentiation among tropical and temperate plants.

## Methods

### Selection of study clades and scoring pollination mode

For selecting study clades, we employed the following criteria: 1) monophyletic groups (regardless of taxonomic level) encompassing both bee and hummingbird pollination, with 2) considerable phylogenetic understanding of the lineage (note that we included all taxa with available pollinator information, regardless of the direction of pollinator shifts), 3) clades occurring across elevational gradients in the Americas, and 4) lineages with empirical pollinator observations, and, additionally, detailed, system-specific assessments of pollination syndromes (floral trait combinations useful for predicting pollinators for species lacking empirical observations, Dellinger 2020). We based our initial searches on lists from Tripp & Manos 2008 and Abrahamczyk & Renner 2015. In addition, we searched for pollinator records for well-known Neotropical clades with shifts to hummingbird pollination (Bromeliaceae, Gesnerioideae, *Iochroma*, Loranthaceae, Merianieae, *Palicourea, Psychotria*) using keyword searches in google scholar (i.e. clade name + pollinat*). This resulted in an initial list of 2563 taxa across 26 groups.

For each taxon, we extracted information on the most efficient pollinator (either empirical or inferred through pollination syndromes) from the respective paper (Table S1, see original dataset deposited in Dryad). We summarized these detailed data into five categories: bee (n=1141), insect (n=121, including butterfly, fly, moth and generalized insect-pollinated systems), hummingbirds (n=768), mixed hummingbird-insect systems (n=115), other vertebrate pollinators (n=87, including bats, mammals and other birds).

### GBIF occurrence data and environmental variables

All following steps were performed in the programming environment R (R Development Core team 2021). We screened the initial plant taxon list (n = 2563) using *Taxonstand* (Cayuela et al. 2021) to correct spelling mistakes and synonyms. Next, we submitted the list to GBIF to search for occurrence data for each taxon (*rgbif*, Chamberlain et al. 2021). We applied strict filtering using the function *occ_issues*, filtering out records with a continent-country mismatch, country-coordinate mismatch, records where the continent classification was derived from coordinates, invalid continents or coordinates, zero coordinates, coordinates out of range, presumed swapped coordinates, invalid geodetic datum, fuzzy taxon matches and non-metric, non-numeric or unlikely elevation (732,047 records). Next, we filtered data using *CoordinateCleaner* (Zizka et al. 2018) removing records located in country centroids, the sea, around gbif headquarters, duplicates or records with equal longitude and latitude (leaving 548,251 records). We then removed records outside of the Americas and calculated median elevation, median latitude and median longitude for each taxon, leaving 1807 taxa. Taking the median per taxon further minimizes bias due to potential single erroneous occurrence points.

Since many taxa had been removed due to missing elevation, we repeated the GBIF query for those, filtering for coordinate-related issues and through *CoordinateCleaner*, but not filtering out problematic elevation records (8913 records). For each of these records, we extracted an elevation value from a global 30 arc sec elevation raster (Amatulli et al. 2018, *raster*, Hijmans 2021) and, again, calculated median elevation, latitude and longitude per taxon. We ran an additional, manual screening of the taxon list to identify misspelled taxon names or synonyms that had not been corrected by *Taxonstand*, and downloaded additional data as specified above. Further, since some taxa may have gotten lost by the strict filtering settings, we ran another GBIF query for the remaining taxa only filtering through CoordinateCleaner (adding five more taxa). This left us with a final dataset of 2232 taxa across 26 study groups across the Americas (Table S1, Fig. S1).

In order to evaluate whether different pollination strategies associate with specific temperatures or levels of precipitation, we downloaded layers for mean annual temperature (bio1) and precipitation (bio12) from the worldclim dataset at 30 arc sec resolution (Hijmans 2017). In addition, we downloaded data on mean annual cloud cover (https://www.earthenv.org/cloud) since high cloud cover strongly impacts flight activity of poikilothermic insects, but not of birds (Cruden 1972). Next, for each pruned occurrence record, we extracted the respective bio1, bio12 and cloud cover value and calculated the median per taxon. The final dataset is available in the online repository Dryad (*link to be included*).

### Ecogeographic modelling

To summarize ecogeographic patterns in our dataset, we plotted occurrence data on a map of the Americas (*maptools*, Bivand & Lewin-Koh 2021). We used Whittaker biomes (Whittaker 1975, Ricklefs 2008, *plotbiomes*, Stefan & Levin 2021) to assess whether vertebrate- or insect-pollinated species in our sample associate with different biomes. We used Chi-square tests to test for differences in biome occupation and used correlation plots (*corrplot*, Wei & Simko 2021) to visualize the relative contribution of Pearson residuals to Chi-squares (Fig. 1).

**Figure 1.**
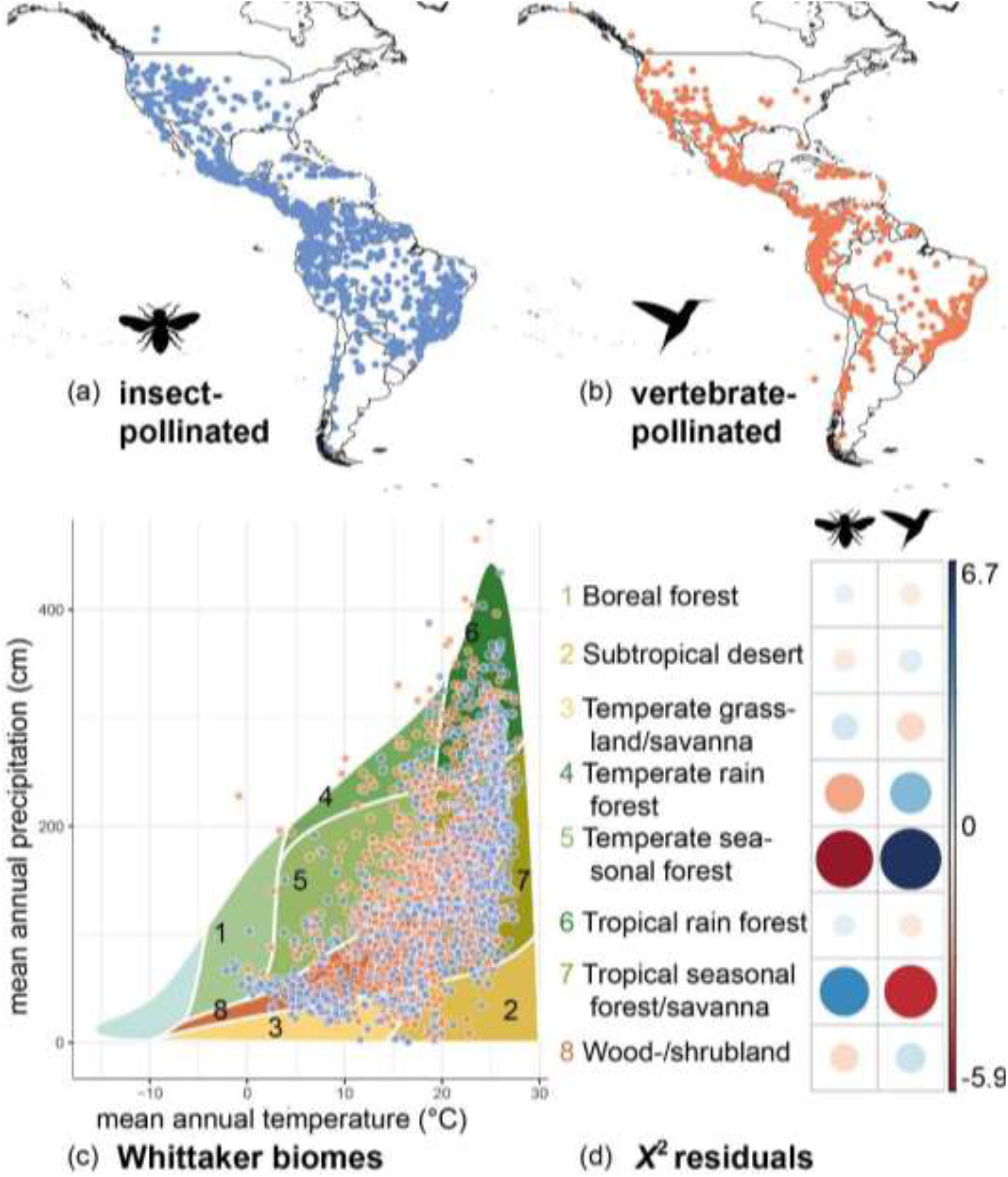
Geographical and biome distribution of insect- and vertebrate-pollinated species included in our dataset. (a) Insect-pollinated species (n = 1262) and (b) vertebrate-pollinated species (n = 970) across the Americas; each dot represents the median occurrence of each species. (c) Vertebrate-pollinated species in our dataset were found primarily under intermediate mean annual temperature (10-20°C) and mean annual precipitation (e.g. in temperate seasonal forests), while insect-pollinated species also occurred under cooler and drier (e.g. in temperate grasslands, shrublands) and warmer (e.g. in tropical seasonal forests) conditions. (d) Contribution of biomes (absolute standardized residuals) to total *X*^2^, positive values (blue) indicate a positive relation, negative values (red) indicate a negative relation, the size of the circle indicates the strength of the relationship.

We used generalized linear mixed effects models (GLMMs, *lme4*, Bates et al. 2015) to test whether associations between pollination mode and elevation or climate depend on latitude. Since elevation was strongly correlated with bio1 and bio12 (Fig. S2), we only included elevation, latitude and cloud cover in our model. Further, we decided to merge the five pollinator groups into a binary response variable (insect versus vertebrate) for two reasons. First, visualizing our data showed similar patterns among species classified as bee or insect pollinated, and among species classified as hummingbird, mixed-hummingbird or other-vertebrate pollinated (Fig. S3). Second, species initially classified as other-vertebrate pollinated were restricted to the tropics, which would bias model estimation at higher latitudes. We then constructed binomial GLMMs (logit link), testing for an interaction of elevation, cloud cover and latitude. We included the respective study clades as a random effects variable, testing models with random intercepts, and random slopes (e.g. elevation|taxonomic group) and intercepts. Since initial models did not converge, we scaled and centered the numeric predictor variables and optimized models using *bobyqa* (Bates et al. 2014) across 2e^5^ iterations. We ran stepwise model simplification (drop1) to identify the most parsimonious model and used Chi-square tests and ANOVA to check whether the simplified model led to a significant reduction in the residual sum of squares. To validate the final model, we visually checked Pearson residuals, predicted versus original values and the ROC (receiver operating characteristic curve, *pROC*, Robin et al. 2011, Fig. S5, S7). We visualized fixed and random effects using *sjPlot* (Lüdecke 2021), *effects* (Fox & Weisberg 2018) and *ggplot2* (Wickham 2016). Finally, to assess potential bias arising from uneven sampling across study groups and pollination strategies, we randomly subsampled our dataset to 25% 100 times, reran model selection, refit the best-fit models and summarized model coefficients.

Given the marked differences detected among the tropics and temperate zone, we split the dataset into ‘tropics’ (−30° - +30° latitude), ‘temperate North’ (> 30° latitude) and ‘temperate South’ (< 30° latitude) to evaluate the effects of mean annual temperature, precipitation and cloud cover on pollination mode. We followed the same approach as outlined above to determine the best-fit model.

While we accounted for some phylogenetic structure in our GLMMs by specifying the study groups as random effects, we additionally ran phylogenetic generalized linear mixed models for more accurate assessment of phylogenetic non-independence of data points. To this end, we downloaded the angiosperm-wide phylogeny provided by Zanne et al. (2014) and Smith & Brown (2018) through Jin & Qian (2019, *V*.*PhyloMaker*) and pruned it to the species included in our sample. Approximately 41% of our study species (932) were covered by this phylogeny, with even sampling across the distribution range and pollination strategies (472 insect-pollinated species, 460 vertebrate-pollinated; Fig. S4, Table S2), hence suitable for investigating large-scale patterns in pollination strategies. On this subset, we built binary phylogenetic GLMMs including elevation, latitude and cloud cover under a Brownian motion model of trait evolution (*binaryPGLMM*-function in *ape*, Paradis & Schliep 2019). We refrained from Bayesian fitting to avoid phylogenetic variance getting trapped around zero (a common problem in binomial GLMMs, Paradis & Schliep 2019). Since cloud cover did not show significant effects, we reduced the model to only include an interaction between elevation and latitude (Table S7).

## Results

### Flipped elevational distribution of insect and vertebrate pollination across latitudes

Our final dataset included 2232 taxa across 26 study groups (22 families) across the Americas (Fig. 1), with 1262 species classified as insect-pollinated and 970 as vertebrate-pollinated. Our results showed that Cruden’s model of vertebrate pollinators dominating at higher elevations only holds in the Tropics (Fig. 2). Here, vertebrate-pollinated species occurred significantly more often at higher elevations (above ca. 1400 meters) than insect-pollinated species. This pattern was reversed at higher latitudes of temperate North and South America where insect-pollinated species occurred at higher elevations (above 1000 – 1500 meters) than vertebrate-pollinated species (Fig. 2). The flipping point between vertebrate or bee pollination at high elevations, which we term the ‘Tropical flip’ of pollination mode, occurred at 27° in the Northern and -30° in the Southern Hemisphere. These patterns were statistically supported when jointly modelling the effect of altitude, latitude and cloud cover on pollination mode (binomial GLMMs using insect pollination as reference level, significant interaction between elevation and latitude, but not cloud cover: *likelihood-ratio 65*.*5*, estimate -0.51, *z-value – 4*.*463, p < 0*.*001, AUC 0*.*807*, Table 1, Fig. 2, Table S4, Fig. S5). These results were consistent when randomly sub-setting the data to 25% (96 of 100 comparisons showed *p < 0*.*01* effects of *elevation*latitude*, Table S5, S6) and when accounting for phylogenetic relatedness (Table S7, Fig. S6). Furthermore, we recovered the Tropical flip in the majority of our individual clades, affirming the generality of this pattern at a broad scale and within lineages (Fig. S7, e.g. tropical: Bromeliaceae, Gesnerioideae, Merianieae, *Passiflora, Psychotria* and *Palicourea*; temperate: *Aquilegia, Castilleja, Penstemon*, whole distribution: *Ruellia, Salvia*). Moreover, tropical lineages which are uncommon at elevations higher than 1400 meters (e.g. *Costus*, Bignonieae) did not show altitudinal differentiation of pollinators.

**Table 1.** Best-fit GLMMs on the effect of elevation, latitude and bioclimatic variables on pollination mode. Models on the full dataset and the tropics included random slopes and intercepts with elevation and temperature, respectively, for the study clades. bio12 – mean annual precipitation; see Table S4 for details on model selection.

| **full-model** | **Estimate** | **std-err** | **z-value** | **p-value** |
| --- | --- | --- | --- | --- |
| (Intercept) | -0.24655 | 0.28717 | -0.859 | 0.391 |
| elevation | 0.83913 | 0.19826 | 4.232 | <0.001 |
| abs(latitude) | 0.08466 | 0.11181 | 0.757 | 0.449 |
| **elevation:abs(latitude)** | **-0.50585** | **0.11335** | **-4.463** | **<0.001** |
| **tropics** |  |  |  |  |
| (Intercept) | 0.94678 | 0.33336 | 2.840 | 0.00451 |
| **cloud cover** | **0.31511** | **0.06725** | **4.685** | **<0.001** |
| **temperate North** |  |  |  |  |
| (Intercept) | -0.9452 | 0.4313 | -2.192 | 0.02841 |
| **bio12** | **0.5996** | **0.2051** | **2.923** | **<0.01** |
| **cloud cover** | **-13.084** | **0.2418** | **-5.412** | **<0.001** |
| **temperate South** |  |  |  |  |
| (Intercept) | 0.1035 | 0.7781 | 0.133 | 0.8942 |
| **bio12** | **14.595** | **0.6877** | **2.122** | **0.0338** |

**Figure 2.**
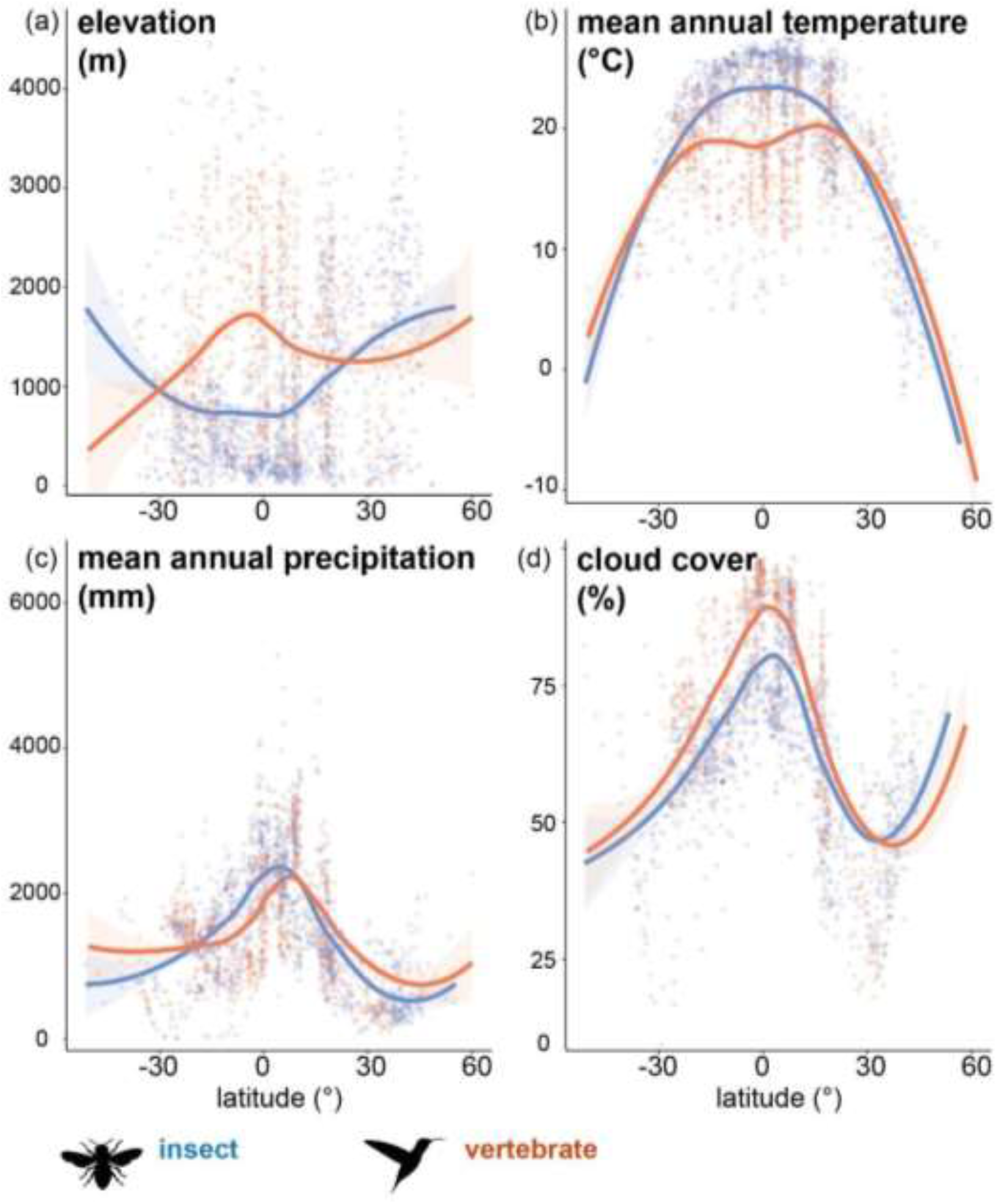
Relation between pollination mode, elevation and different bioclimatic variables across latitudes. (a) Marked differentiation in the distribution of insect- and vertebrate-pollinated species across latitudes, with vertebrate-pollinated species occurring at higher elevations than insect-pollinated species in the tropics, but not in temperate regions. (b) Vertebrate-pollinated species associate with lower mean annual temperatures in tropical regions than insect-pollinated species, but with slightly warmer conditions in temperate zones. (c) Variable relationships among pollination strategies and mean annual precipitation, with bee-pollinated species in our dataset occurring under slightly wetter conditions than vertebrate-pollinated species around the equator. (d) Vertebrate-pollinated species occur in areas with higher cloud cover than bee-pollinated species in the temperate South and the tropics.

### Vertebrate pollination associated with intermediate temperatures and moist conditions across latitudes

Cruden (1972) experimentally showed that bee-pollinator efficiency was reduced under cool, cloudy and rainy conditions. We hence tested whether abiotic climatic variables (mean annual temperature, precipitation and cloud cover) consistently associate with either insect or vertebrate pollination (Fig. 2b-d). To avoid confounding effects driven by the tropical flip’, we split the dataset into tropical (grossly defined as +/- 30°) and temperate zones for statistical analyses. Overall, mean annual temperature was strongly positively correlated with elevation (Fig. S4). We found vertebrate-pollinated species occur at lower temperatures than insect-pollinated species in the tropics (ca. 18°C versus 23°C, Fig. 2b), but did not find a clear association in temperate zones. Temperature was not significant in any of the models and never retained in the best-fit models (Table 1, S4). Among tropical species, vertebrate pollination most strongly associated with high levels of cloud cover (Table 1, Fig. 3b), with the best-fit model allowing for random slopes and intercepts with temperature for the different clades (Table 1, S4, *variance 1*.*66, correlation 0*.*88; AUC 0*.*824*, Fig. S8). Among temperate North American species, we found significant additive effects of cloud cover and precipitation (Fig. 3c), with vertebrate-pollinated plants more likely to occur in areas with low cloud cover but high precipitation (*AUC 0*.*830*, Table 1, S4, Fig. S8). In temperate South American species, vertebrate-pollinated species again showed significant associations with increased precipitation but not cloud cover (*AUC 0*.*928*, Table 1, S4, Fig. 3d, Fig. S8).

**Figure 3.**
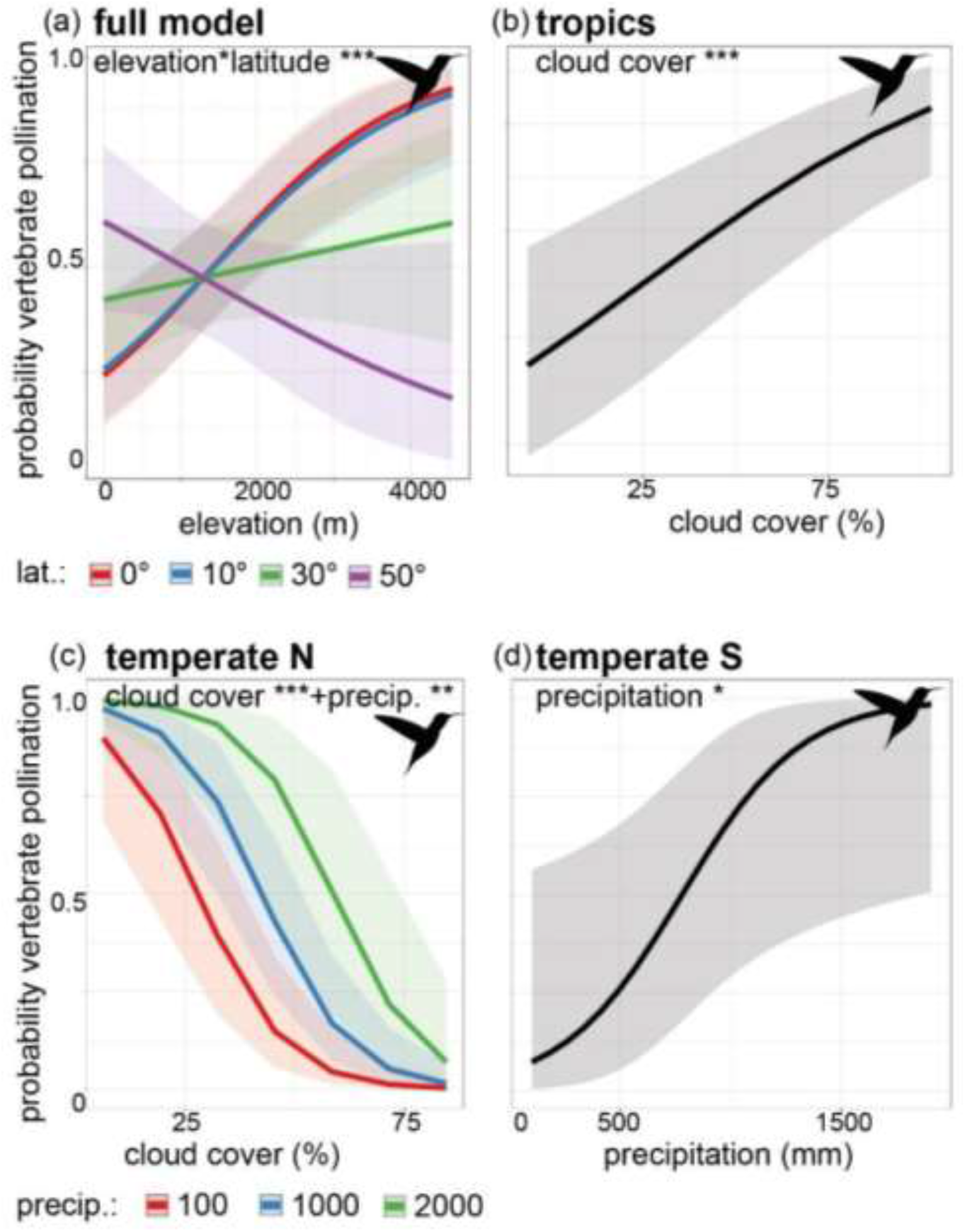
Results of the best-fit models investigating the effects of different bioclimatic variables across latitudes. (a) The probability of vertebrate-pollination increases significantly across elevation in the tropics (0°, 10° latitude) while it gradually decreases towards higher latitudes (30°, 50°). (b) The probability of vertebrate-pollination increases significantly with increased cloud cover in the tropics (i.e. montane cloud forests, n = 1917). (c) In the temperate zones of the Northern hemisphere, the probability of vertebrate pollination decreased significantly with increasing cloud cover, but less so when precipitation was high (n = 265). (d) In the temperate zones of the Southern hemisphere, the probability of vertebrate pollination increased with increasing precipitation (n = 50). Insect pollination was used as reference level in the binomial GLMMs (and is hence not depicted), 95% confidence intervals are given, *** p < 0.001, ** p < 0.01, * p < 0.05.

Finally, since the abiotic climatic environment structures the distribution of major vegetation types and animal groups on earth, we tested whether insect and vertebrate pollination associate with particular broad-scale biogeographic biomes (Whittaker 1975). We sorted species into biomes based on temperature and precipitation and found significant differences in biome occupation between insect- and vertebrate-pollinated species, mainly driven by temperature (*X*^*2*^ *= 149*.*81, df = 7, p-value < 0*.*001*, Table S3, Fig. 1). While both pollination strategies were found in each biome (except Tundra), vertebrate-pollinated species were significantly more common at intermediate temperatures (10°C-20°C) and precipitations in temperate seasonal forests and temperate rainforests (Fig. 1, Table S3). Insect-pollinated species occurred significantly more often under warmer conditions in tropical seasonal forests/savannas, and under both cooler and drier conditions in wood-/shrublands, boreal forests and temperate grasslands (Fig. 1, Table S3).

## Discussion

We here present fundamental differences in the altitudinal distribution of insect- and vertebrate-pollinated plant species across the Americas, rejecting the hypothesis that shifts between insect and vertebrate pollinators occur universally along altitudinal gradients (Cruden 1972). We find, however, consistent associations between pollination mode and critical climatic variables, which disproportionately impact the activity of ectothermic insects, but not of endothermic vertebrates. We recover this pattern across multiple. plant clades, underscoring its generality. Our results affirm that shifts among insect and vertebrate pollination have been repeatedly influenced by climate-related alterations in pollination efficiency (Cruden 1972, Thomson & Wilson 2008). In the following, we discuss our findings in light of macroevolutionary and macroecological differences among tropical and temperate zones in the Americas.

Building on Cruden’s (1972) model of reduced pollination efficiency of ectothermic bees under adverse weather conditions in tropical mountains, one major open question is why hummingbird-pollinated plants do not commonly occur at high elevations also in the temperate Americas. There are several possible explanations to this pattern. In Western North America, the distribution of hummingbird pollination is shaped by seasonal climatic conditions and migratory patterns of hummingbirds. Flowering windows are constrained by (late) snow melt at high elevations and summer drought at lower elevations, causing marked seasonal variation in nectar resources across elevation. During their summer breeding seasons, migratory hummingbirds spend extended time at low- to mid-elevations below the treeline since they depend on the close availability of abundant nectar resources and nesting sites (i.e. trees, which are absent above the treeline; Grant & Grant 1968, Stebbins 1989). Higher elevations in temperate North America, on the other hand, are characterized by exceptionally diverse bumblebee communities (29 species occur at high elevations, McCabe & Cobbs 2021). Bumblebees are, in contrast to most other bees, effective cold-adapted high elevation pollinators (McCallum et al. 2013). Bumblebees may hence start visiting flowers right at the start of the flowering season (when hummingbirds are still migrating or already nesting), and possibly effectively occupy the available pollination niche space throughout the short high-elevation flowering periods (Pyke et al. 2011). In tropical mountains, the ‘bumblebee’ pollination niche space seems less saturated since comparatively few large, cold-adapted bee species (e.g. from the genera *Bombus, Centris, Eufriesia, Eulaema, Xylocopa*) reach the montane cloud forest or Páramo zones (Gonzalez & Engel 2004).

The Tropical flip of insect and vertebrate pollination across elevation is particularly interesting in light of contrasting diversification histories between tropical and temperate plant lineages. While evolutionary shifts to vertebrate (and particularly hummingbird) pollination correlate with large radiations in numerous tropical clades, shifts to hummingbird pollination in temperate zones are often restricted to single species (Grant & Grant 1968, Abrahamczyk & Renner 2015, Wessinger et al. 2019). Focusing on the intriguing tropical radiations, some authors have hypothesized that hummingbird pollination itself increases diversification rates (e.g. Serrano-Serrano et al. 2017). This claim has recently been criticized as too simplistic and disconnected from potential underlying mechanisms (Kessler et al. 2020). Our results support this criticism and suggest that, instead (or in addition), the montane environment that hummingbird/vertebrate pollination is often linked to in the tropics (but not in temperate zones), may drive the observed tropical radiations (Lagomarsino et al. 2016). In fact, rugged mountain topography, creating exceptional microclimatic variation across small spatial scales, and hence potential for allopatric divergence, is usually regarded as a key ingredient for evolutionary radiations. This idea is supported by the fact that diversification rates are comparably high in animal-pollinated plants and tropical ferns (which are not pollinated by animals, Testo et al. 2018). In addition, tropical hummingbirds are functionally more diverse (i.e. different bill lengths/shapes), possibly resulting in a more fine-grained subdivision of the hummingbird pollination niche in the tropics. Overall, however, it remains largely unexplored whether early evolutionary shifts to vertebrate pollination enabled plant lineages to colonize tropical mountains, where they then diversified within the vertebrate pollination niche, or whether adaptive abiotic niche evolution into tropical mountains preceded shifts to vertebrate pollination. These questions may best be addressed in clade-specific studies by combining refined niche-modelling with detailed phylogenetic analyses.

The lack of studies jointly analyzing abiotic and biotic processes in plant diversification is quite surprising given that the classical pollinator shift model establishes a strong link between the two. This model posits that externally (e.g. climate) induced changes in the abundance and hence efficiency of a plant species’ ancestral pollinator may trigger shifts in pollination mode (Thomson & Wilson 2008). Our finding that humidity-related factors (precipitation, cloud cover), along with temperature, differentiate insect- and vertebrate-pollinated plants both in the tropics and temperate zones, supports this concept (Grant & Grant 1968, Krömer et al. 2006, Chalcoff et al. 2012). Shady, moist, forested habitats (which applies to both tropical cloud forests and temperate wooded ravines at lower elevations) may indeed represent “hummingbird habitats” (Stebbins 1989). These habitats may feature pollination niche space that is unoccupied by bees, and hence filled by vertebrates (Stebbins 1989). Thus, our results support the notion that evolutionary shifts in pollination strategies are intimately linked to the occupation of different abiotic niche space. Pollinator shifts are, however, not always linked to abiotic niche differentiation, since, for example, both insect- and vertebrate-pollinated plants occur in lowland rainforests (Fig. 2). In these cases, shifts among insect and vertebrate pollination must, consequently, result from factors other than climate-related alterations in pollination efficiency. Spatial and temporal variation in food availability, along with competition among pollinators, may drive such shifts (Abrahamczyk et al. 2011, Temeles et al. 2016).

In addition to abiotic climatic factors, there are fundamental differences in the build-up and complexity of plant-pollinator interactions across latitudes, which should be considered in the interpretation of our results. First, the diversity of vertebrate pollinators in the tropical Americas (i.e. hummingbirds, passerines, bats, rodents, lizards) is unparalleled in temperate zones (only 26 hummingbird species pollinate at least 240 temperate plant species, Abrahamczyk & Renner 2015). While we did not analyze these vertebrate pollinator groups separately given the relative scarcity of data, we found a strong trend for species pollinated by bats, rodents or passerine birds to occur at the highest elevations in the tropics (Fig. S3). In addition, several of our study clades (i.e. Bromeliaceae, centropogonids, Merianieae), exhibit stable “bimodal” pollination systems. In these systems, the pollination niche is effectively partitioned among diurnal (i.e. hummingbirds) and nocturnal (i.e. bats) pollinators (Dellinger et al. 2019, Lagomarsino & Muchhala 2019). The 24/7 pollination services these flowers receive may be particularly adaptive in the tropical montane forests by buffering reductions in flower visitation frequency in periods of heavy rainfall. No nocturnal vertebrate pollinators are available in the temperate zones, however, where hawkmoths constitute the only specialized nighttime pollinators. Hawkmoths are, however, like bees, ectothermic and hence restricted in their flight capacity to warm conditions (Cruden 1976).

Second, pollination systems in the tropics are generally found to be more specialized than in temperate zones, with high competition among pollinators driving fine-grained partitioning of the available pollination-niche space (Trojelsgaard & Oleson 2013). In connection with this, among-pollinator competition has been proposed as the driver behind the elevational distribution of moth pollination (Cruden 1976). In fact, moth pollination shows a similar Tropical flip as observed in our study, and is hence worth considering here: hawkmoths act as effective pollinators in (warm) tropical lowlands, but pollinate at (cooler) high elevations in the temperate zone. The ability to act as effective pollinators in temperate mountains is brought about by a behavioral adaptation to be active during warmer daytime hours (Cruden et al. 1976). In tropical mountains, on the other hand, the pollination niche seems effectively saturated by hummingbirds, and competition for pollination niche space may hence restrict hawkmoths to nocturnal pollination and lowland rainforests there (Cruden et al. 1976). Several of the temperate clades included in our study also encompass species pollinated by hawkmoths (e.g. *Aquilegia, Castilleja, Ipomopsis, Schizanthus, Silene*). While we had to lump hawkmoth pollinators with other non-bee insect pollinators due to the small sample size, we recovered the same pattern as Cruden (Fig. S3). Taken together, these results suggest a potential large-scale difference in the structure (i.e. competition for/among pollinators) and function (i.e. fitness benefits or trade-offs related to pollinator generalization) of plant-pollinator interactions between the tropics and temperate zones.

Finally, we want to highlight the particular need for more research on insect pollination in a macroevolutionary context particularly in temperate zones. Although only about 1% of temperate North American plant species are hummingbird pollinated (Abrahamczyk & Renner 2015), these have been much more researched than related insect-pollinated species, leaving painfully large gaps in our understanding of the ecology and evolution of insect pollination. Further, the traditional narrow focus on hummingbird-bee sister-species pairs in temperate North America limited our capacity of reliably determining related bee-/insect-pollinated species in many lineages where hummingbird pollination evolved. In tropical lineages, where taxonomic relationships often remain more elusive (and sister-species pairs can hence not be identified reliably), broad-scale, comparative analyses, using pollination syndrome approaches are more common, allowing for the inclusion of more tropical lineages in the present study. Overall, broadening the scope of pollination studies by addressing plant-pollinator interactions as functions of their abiotic, but also biotic, environment both at the scale of interacting communities and evolving entities (i.e. clades) is needed in the close future, not only for gaining a more profound perspective on the ecogeographic processes structuring i.e. the Tropical flip, but also in light of current, climate-driven changes in plant-pollinator interactions (Franzén & Öckinger 2012).

## Acknowledgements

We thank the labs of Stacey Smith and Erin Manzitto-Tripp for inspiring discussions on this topic and helpful feedback on the project. We particularly thank Miranda Sinnott-Armstrong for support with and recommendations for data analyses. We further thank Andreas Berger for sharing data on pollination in Rubiaceae with us. This work was supported by FWF-grant T-1186 to ASD, NSF DEB-2052904 to CAW, and NSF-1553114 to SDS.

## Data availability statement

All data analyzed in the manuscript will be made accessible through the public Dryad repository.

## Author Contribution

All authors conceived the study, ASD and AH compiled the species datasets, ASD ran the analyses and drafted the manuscript, AH, CW and SDS contributed to improving and revising the manuscript.

## Biosketch

Agnes Dellinger is interested in investigating the relative importance of abiotic and biotic processes in driving plant diversification at macro- and microevolutionary scales, and how flowers diversify and evolve under continuous or divergent pollinator-mediated selection. She mostly works on the pantropical plant family Melastomataceae. Find more about her research here: https://agnesdellinger.org/

