## Supplemental Information for "Flipped elevational pattern of pollination mode in tropical vs. temperate Americas"

**Supplementary Information for Flipped elevational pattern of pollination mode in tropical vs. temperate  America**

**Table S1:** Plant clades, number of plant taxa per pollinator group, total number of taxa considered and clade sizes of focal study clades. Please note that the ratio observed/syndrome may not reflect the absolute number of available empirical pollinator observations per clade (e.g. *Salvia*) but is based on the respective data source used in our study. We here highlight the most important literature sources for pollinator information per clade, more details may be found in our dataset deposited in Dryad.

| Family | study group | insect | bee | humming bird | mixed humming bird | other vertebrate | total | observed/ syndrome | clade size | literature source |
|---|---|---|---|---|---|---|---|---|---|---|
| Acanthaceae | *Ruellia* | 7 | 65 | 40 | 4 | 4 | 120 | 31/89 | 275 (NW) | Tripp & Manos 2008; Tripp & Tsai 2017 |
| Bignoniaceae | Bignonieae | 33 | 304 | 21 | 5 | 3 | 366 | 42/324 | 393 | Alcantara & Lohmann 2010 |
| Bromeliaceae | Bromeliaceae | 2 | 42 | 62 | 16 | 31 | 153 | 39/114 | 3140 | Aguilar-Rodríguez et al. 2019; Givinish et al. 2014 |
| Campanulaceae | Centropogonids | 0 | 5 | 14 | 0 | 11 | 30 | 5/25 | 550 | Lagomarsino et al. 2017 |
| Caryophyllaceae | *Silene* | 5 | 2 | 6 | 0 | 0 | 13 | 8/5 | 70 (NAm) | Abrahamczyk & Renner2015 |
| Convolvulaceae | *Ipomoea* | 5 | 12 | 2 | 0 | 2 | 21 | 21/0 | 650 | Rosas-Guerrero et al. 2010 |
| Costaceae | *Costus* | 0 | 19 | 20 | 0 | 0 | 39 | 11/28 | 57 (NW) | Vargas et al. 2020 |
| Gentianaceae | Gentianella | 9 | 0 | 2 | 0 | 0 | 11 | 0/11 | ca. 200 | von Hagen & Kadereit 2001 |
| Gesneriaceae | Gesnerioideae | 10 | 189 | 318 | 0 | 9 | 526 | 112/414 | 1200 | Serrano et al. 2017 |
| Iricaceae | *Iris* (sect. *Hexagonae*, *Longipetalae*) | 0 | 3 | 2 | 0 | 0 | 5 | 5/0 | 8 | Emms & Arnol 2003 |
| Lamiaceae | *Salvia* | 0 | 186 | 117 | 38 | 0 | 341 | 0/341 | 1000 | Wester & Claßen-Bockoff 2011; Kriebel et al. 2020 |
| Loasaceae | Loasoideae | 0 | 23 | 10 | 6 | 2 | 41 | 27/14 | 200 | Ackermann & Weigend 2006 |
| Loranthaceae | Loranthaceae clade D/E | 6 | 1 | 5 | 7 | 0 | 19 | 12/7 | 248 | Amico et al. 2007 |
| Melastomataceae | Merianieae | 0 | 93 | 0 | 20 | 22 | 135 | 20/115 | 303 | Dellinger et al. 2021 |
| Orobanchaceae | *Castilleja* | 2 | 11 | 31 | 2 | 0 | 46 | 20/26 | 180 | Grant 1994; Chuang & Heckard 1992 |
| Passifloraceae | subgen. *Passiflora* | 0 | 24 | 41 | 0 | 3 | 68 | 10/58 | 250 | Abrahamczyk et al. 2014; Pérez & d'Eckenbrugge 2017 |

| | | | | | | | | | | |
|---|---|---|---|---|---|---|---|---|---|---|
| Phrymaceae | *Mimulus* | 1 | 3 | 3 | 2 | 0 | 9 | 9/0 | 120 | Grant 1994; Karron et al. 1995 |
| Plantaginaceae | *Ourisia* | 7 | 0 | 4 | 0 | 0 | 11 | 0/11 | 33 | Meudt & Simpson 2007 |
| Plantaginaceae | *Penstemon & Keckiella* | 0 | 101 | 27 | 4 | 0 | 132 | 73/59 | 292 | Wilson et al. 2007 |
| Polemoniaceae | *Gilia & Saltugilia* | 1 | 3 | 2 | 0 | 0 | 6 | 6/0 | 72 | Grant & Grant 1965 |
| Polemoniaceae | *Ipomopsis* | 6 | 3 | 4 | 0 | 0 | 13 | 13/0 | 28 | Grant & Grant 1965 |
| Ranunculaceae | *Aquilegia* | 6 | 5 | 8 | 2 | 0 | 21 | 2/19 | 25 | Whittal & Hodges 2007 |
| Ranunculaceae | *Delphinium* (sect. *Diedropetal)* | 0 | 2 | 2 | 1 | 0 | 5 | 5/0 | 67 | Grant 1994 |
| Rubiaceae | *Psychotria & Palicourea* | 18 | 36 | 15 | 6 | 0 | 75 | 75/0 | 1780 | Ferreira et al. 2015; Castro et al. 2004; Sakai & Wright 2008; Mesquita-Neto et al. 2018 |
| Solanaceae | *Iochroma* | 0 | 1 | 11 | 2 | 0 | 14 | 14/0 | 35 | Smith et al. 2008 |
| Solanaceae | *Schizanthus* | 3 | 8 | 1 | 0 | 0 | 12 | 9/3 | 12 | Pérez et al. 2006 |
| | ***sum*** | **121** | **1141** | **768** | **115** | **87** | **2232** | **569/1663** | | |

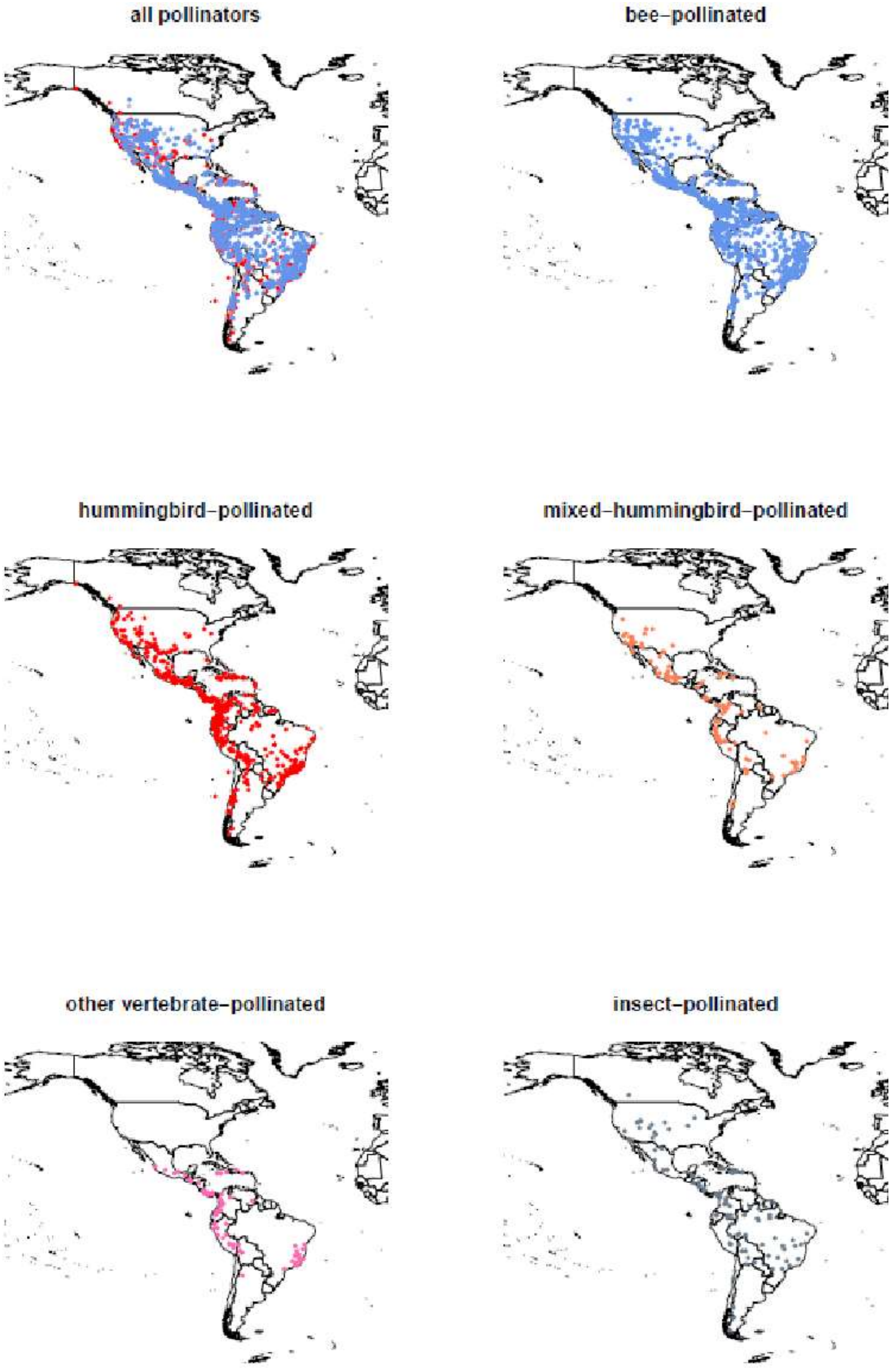


**Figure S1. Distribution of study taxa** in the Americas according to the five pollinator groups.

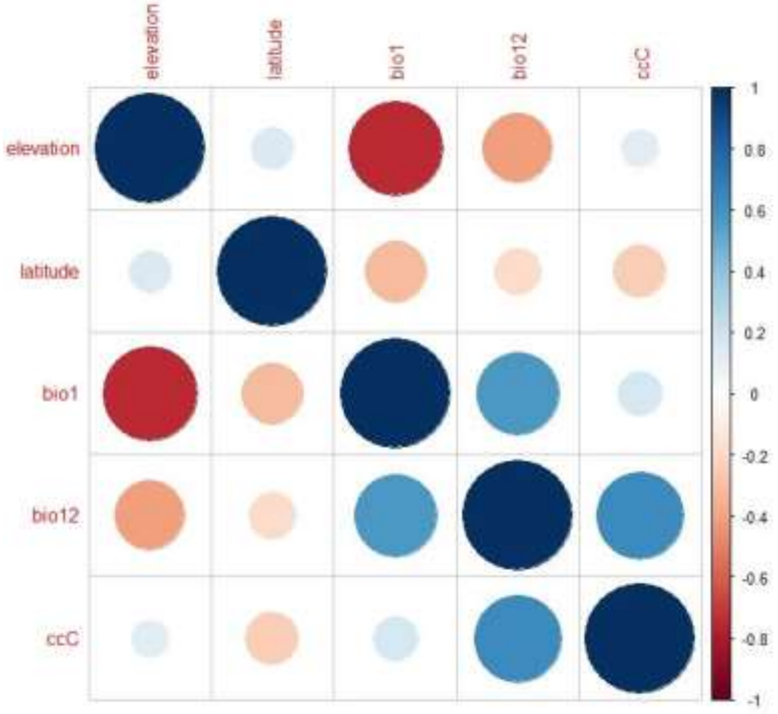


**Figure S2. Correlation structure among our five putative explanatory variables**; elevation, latitude and cloud cover (ccC) were chosen for models run on the whole dataset, while bio1, bio12 and cloud cover were chosen for models on subsets for the tropics and temperate zones.

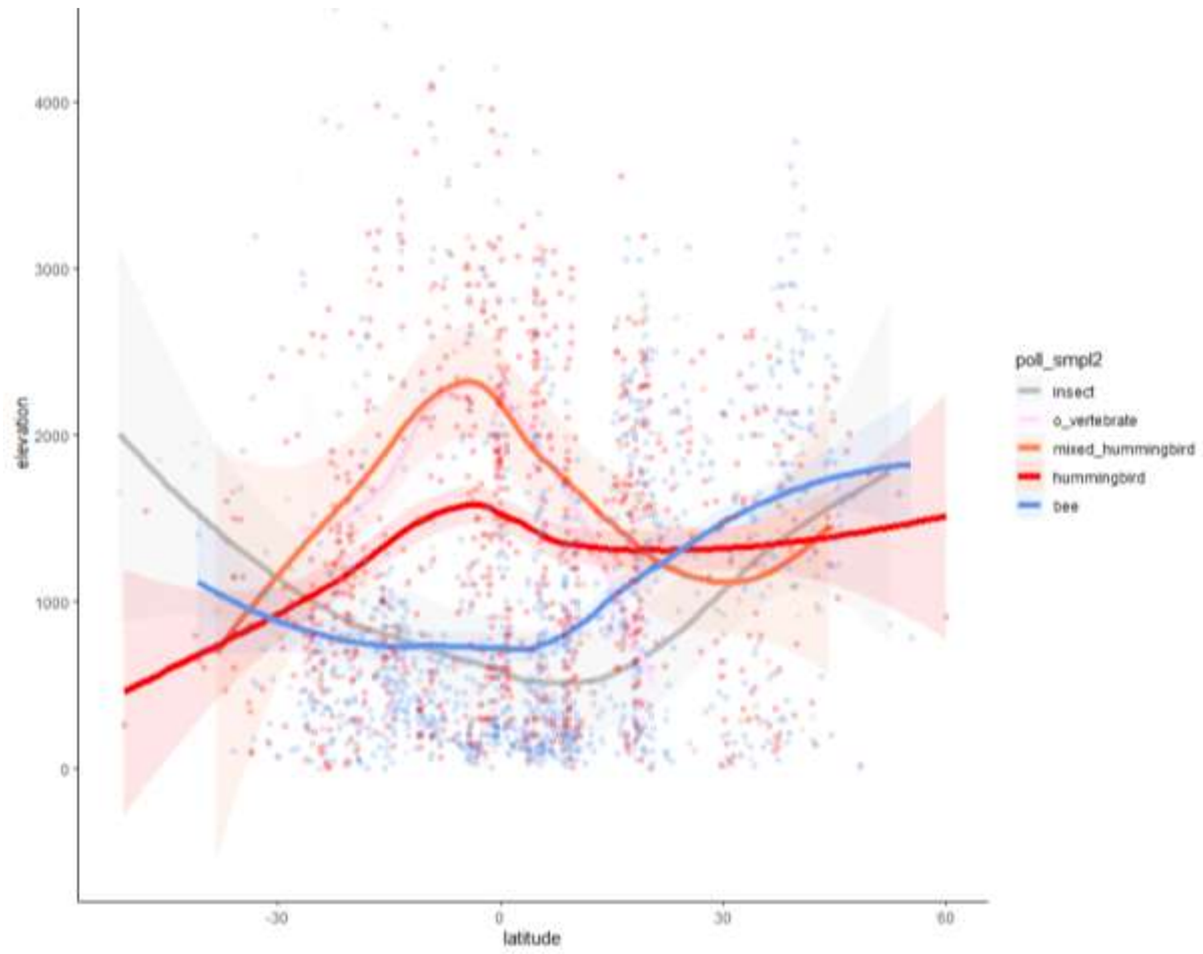


**Figure S3: Elevational and latitudinal distribution of study taxa according to pollination strategy**. Note the similarity in estimated curves among bee and insect pollination, and among hummingbird, mixed-hummingbird and other-vertebrate pollination. We hence merged these groups into either insect or vertebrate pollinated for subsequent GLMM analyses.

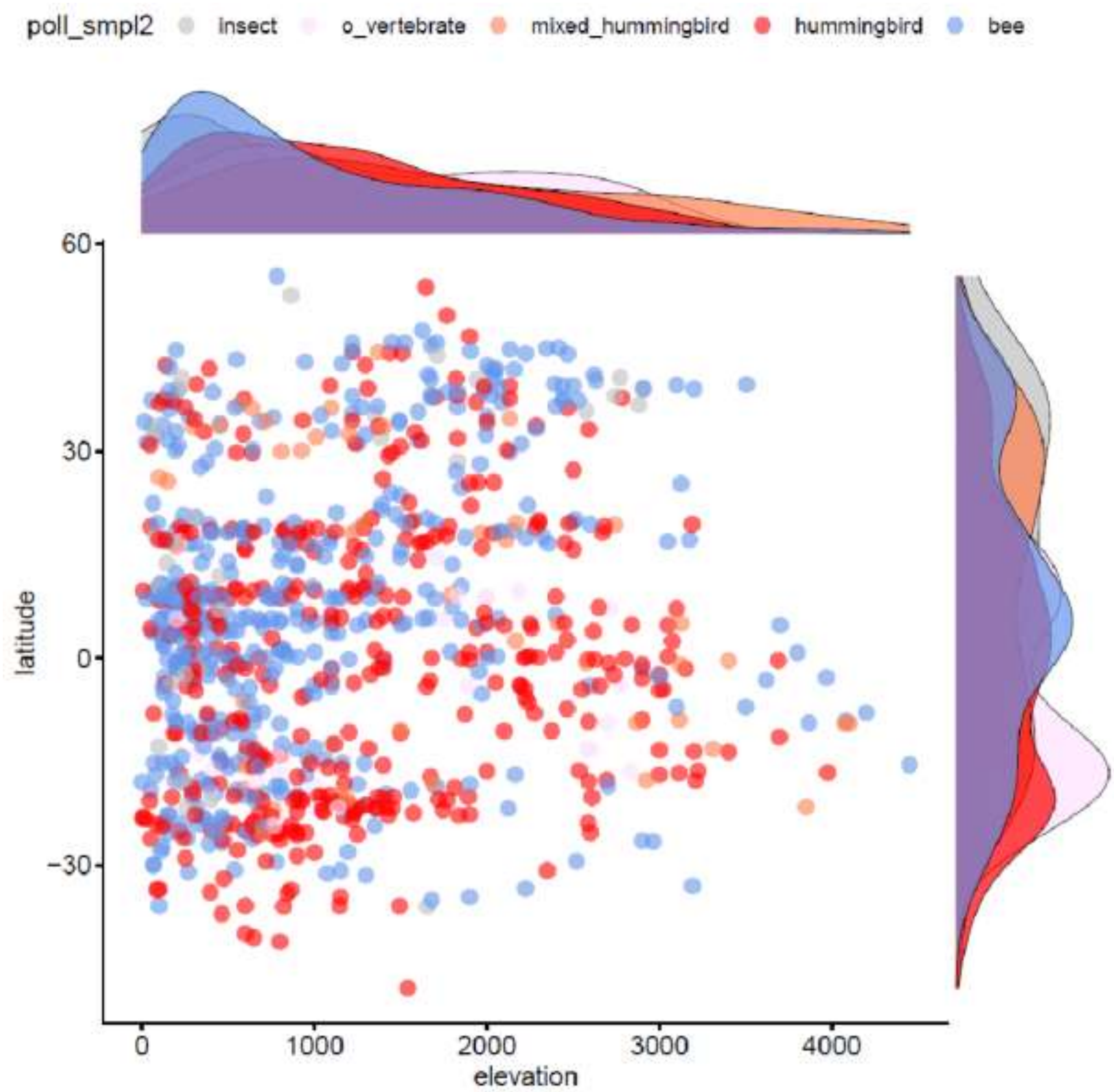


**Figure S4**: Latitudinal and elevational distribution of the 932 taxa represented in the phylogeny; distributions patterns are highly similar to the full dataset (including 2232 taxa).

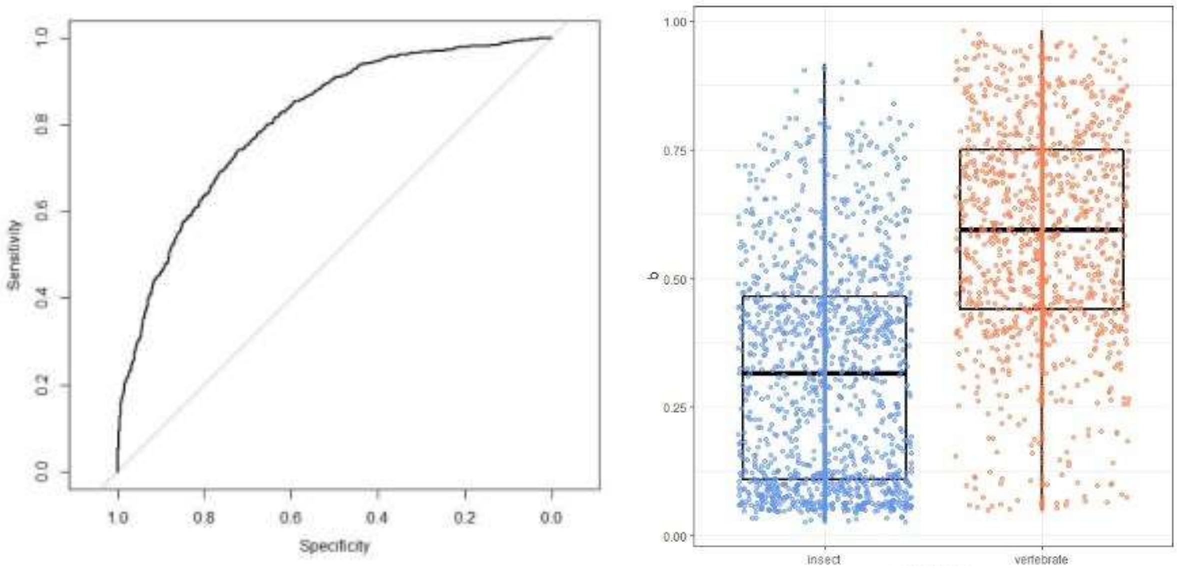


**Figure S5.** ROC best-fit-model on the full dataset (AUC= 0.807) and model predictions for either insect or vertebrate pollination.

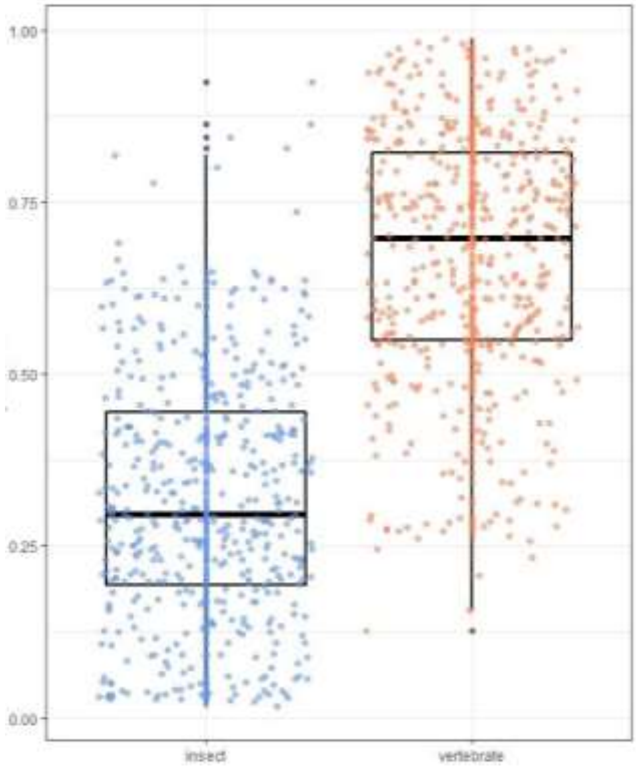


**Figure S6. Predictions for either insect or vertebrate pollination for the 932 species included in the molecular phylogeny.**

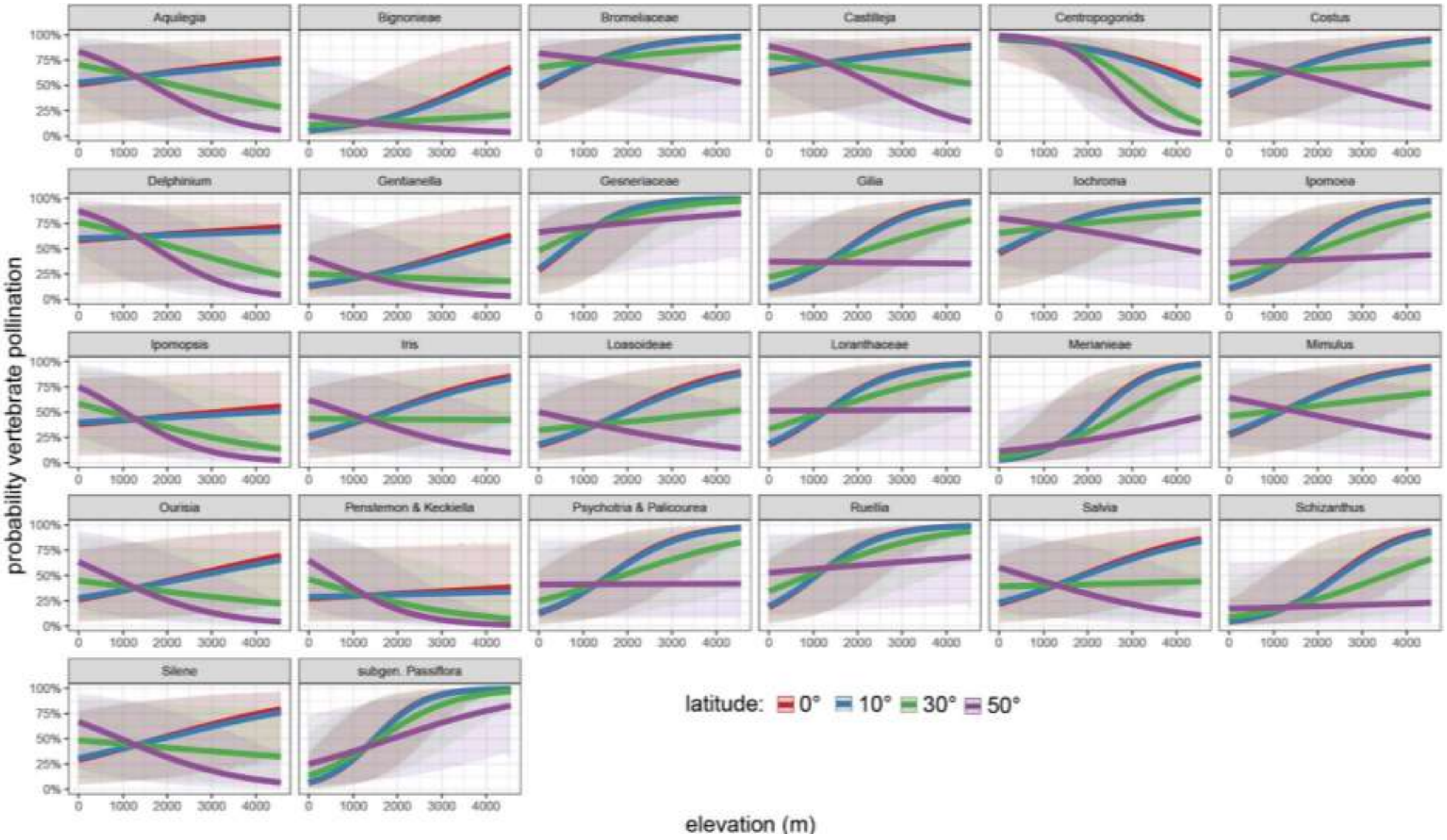
Aquilegia
Bignonieae
Bromeliaceae
Castilleja
Centropogonids
Costus
Delphinium
Gentianella
Gesneriaceae
Gilia
Iochroma
Ipomoea
Ipomopsis
Iris
Loasoideae
Loranthaceae
Merianieae
Mimulus
Ourisia
Penstemon & Keckiella
Psychotria & Palicourea
Ruellia
Salvia
Schizanthus
Silene
subgen. Passiflora
probability vertebrate pollination
0% 25% 50% 75% 100%
0 1000 2000 3000 4000
latitude: 0° 10° 30° 50°
elevation (m)

**Fig. S7. Effects of elevation and latitude on the probability of vertebrate pollination for the different study groups** (included as random effects in our best-fit model (Table S4). Vertebrate pollination is significantly more likely to occur at high elevations in the tropics (0°, 10°), while at high latitudes (50°), vertebrate pollination is found at intermediate to low elevations.

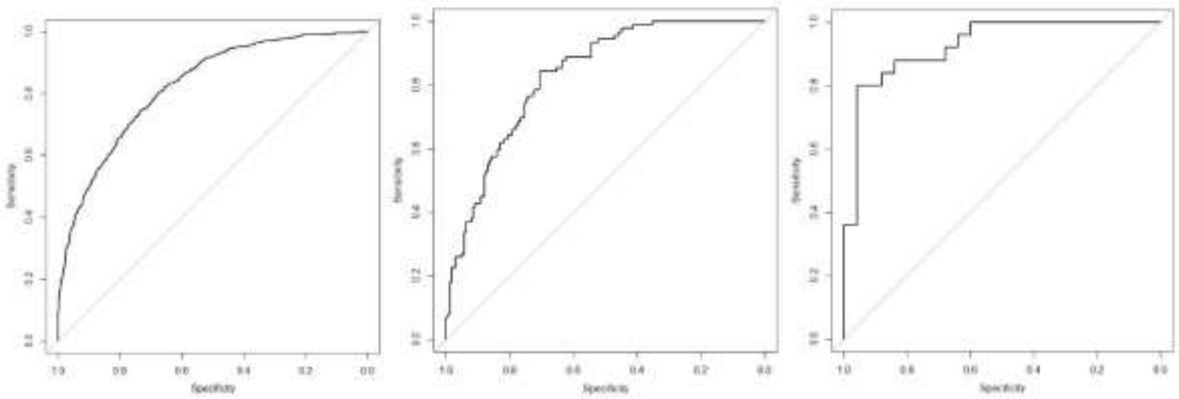

**Fig. S8.** ROC best-fit model on the tropical, temperate North and temperate South dataset.

**Table S2. Proportional representation of the study groups (Table S1) in the molecular phylogeny** (also see Fig. S4 for latitudinal representation). 472 species are insect pollinated (37.4% of the original dataset) and 460 are vertebrate-pollinated (47.4% of the original dataset).

| Taxonomic group | % covered |
|---|---|
| *Aquilegia* | 0.67 |
| Bignonieae | 0.08 |
| Bromeliaceae | 0.71 |
| *Castilleja* | 0.37 |
| Centropogonids | 0.17 |
| *Costus* | 0.87 |
| *Delphinium* | 1 |
| *Gentianella* | 0.09 |
| Gesneriaceae | 0.49 |
| *Gilia* | 0.67 |
| *Iochroma* | 0.86 |
| *Ipomoea* | 0.29 |
| *Ipomopsis* | 0.92 |
| *Iris* | 0.60 |
| Loasoideae | 0.46 |
| Loranthaceae | 0.74 |
| Merianieae | 0.39 |
| *Mimulus* | 0.22 |
| *Ourisia* | 0.18 |
| *Penstemon & Keckiella* | 0.56 |
| *Psychotria & Palicourea* | 0.15 |
| *Ruellia* | 0.63 |
| *Salvia* | 0.31 |
| *Schizanthus* | 0.33 |
| *Silene* | 0.85 |
| subgen. *Passiflora* | 0.74 |

**Table S3. Number of species per Whittaker biome type divided into insect- or vertebrate-pollinated**; the first value gives the actual number of species, the second value gives the number of expected species were there no differences among biomes.

| | insect | vertebrate |
|---|---|---|
| **Boreal forest** | 11 /9 | 5 /7 |
| **Subtropical desert** | 31 /36 | 32 /27 |
| **Temperate grassland/desert** | 66 /57 | 35 /44 |
| **Temperate rain forest** | 17 /32 | 39 /24 |
| **Temperate seasonal forest** | 132 /219 | 254 /167 |
| **Tropical rain forest** | 166 /157 | 110 /119 |
| **Tropical seasonal forest/savanna** | 670 /566 | 329 /433 |
| **Woodland/shrubland** | 163 /180 | 155 /138 |

**Table S4. Summary on model selection for the full dataset and the tropics/temperate zones separately**; pollination strategy was the binary response variable (insect/vertebrate), the best-fit model is highlighted in bolt for each dataset. Bio1 – mean annual temperature, bio12 – mean annual precipitation.

| full dataset (n = 2232) | npar | AIC | BIC | logLik | deviance | Chisq | Df | p-value |
|---|---|---|---|---|---|---|---|---|
| elevation*abs(latitude)+(1\|grouping) | 5 | 2516.2 | 2544.7 | -1253.1 | 2506.2 | | | |
| **elevation*abs(latitude)+(elevation\|grouping)** | **7** | **2487.2** | **2527.2** | **-1236.6** | **2473.2** | **32.962** | **2** | **<0.001** |
| elevation*abs(latitude)+(elevation:abs(latitude)\|grouping) | 7 | 2503.4 | 2543.4 | -1244.7 | 2489.4 | 0.000 | 0 | |
| elevation*abs(latitude)*cloud_cover+(1\|grouping) | 9 | 2517.4 | 2568.8 | -1249.7 | 2499.4 | 0.000 | 2 | 1 |

| Tropics (-30° to +30°; n = 1917) | | | | | | | | |
|---|---|---|---|---|---|---|---|---|
| bio1+cloud_cover+(1\|grouping) | 4 | 2136.1 | 2158.3 | -1064.0 | 2128.1 | | | |
| bio1+bio12+cloud_cover+(1\|grouping) | 5 | 2135.3 | 2163.1 | -1062.6 | 2125.3 | 27.643 | 1 | 0.09639 |
| **cloud_cover+(bio1\|grouping)** | **5** | **2067.3** | **2095.1** | **-1028.6** | **2057.3** | **680.22** | **0** | |
| bio1+cloud_cover+(bio1\|grouping) | 6 | 2068.9 | 2102.2 | -1028.4 | 2056.9 | 0.3874 | 1 | 0.53365 |
| bio1*bio12*cloud_cover+(1\|grouping) | 9 | 2131.9 | 2181.9 | -1057 | 2113.9 | 0.000 | 3 | 1 |

| temperate North (n = 265) | | | | | | | | |
|---|---|---|---|---|---|---|---|---|
| **bio12+cloud_cover+(1\|grouping)** | **4** | **287.50** | **301.82** | **-139.75** | **279.50** | | | |
| bio12+cloud_cover+(bio12\|grouping) | 6 | 287.78 | 309.26 | -137.89 | 275.78 | 37.218 | 2 | 0.1555 |
| bio12+cloud_cover+(could_cover\|grouping) | 6 | 285.15 | 306.63 | -136.57 | 273.15 | 26.300 | 0 | |
| bio1*bio12*cloud_cover*(1\|grouping) | 9 | 287.08 | 319.29 | -134,54 | 269.08 | 40.733 | 3 | 0.2537 |

| temperate South (n = 50) | | | | | | | | |
|---|---|---|---|---|---|---|---|---|
| **bio12+(1\|grouping)** | **3** | **58.183** | **63.919** | **-26.091** | **52.183** | | | |
| bio12+cloud_cover+(1\|grouping) | 4 | 60.024 | 67.672 | -26.012 | 52.024 | 0.1585 | 1 | 0.6905 |
| bio1+bio12+cloud_cover+(1\|grouping) | 5 | 60.726 | 70.286 | -25.363 | 50.726 | 12.983 | 1 | 0.2545 |
| bio1*bio12*cloud_cover+(1\|grouping) | 9 | 65.341 | 82.549 | -23.670 | 47.341 | 33.851 | 4 | 0.4956 |

**Table S5. Summary of model selection for the 25% subset** of the full dataset (558 species); we ran 100 random subsets and determined the best-fit model based on lowest AIC; we present average AIC values across 100 subsets here. Model 1 and 2 resulted as best fit most of the time.

| 100 random subsets to 25% | no of parameters | no times best fit | average AIC |
|---|---|---|---|
| **1. elevation*abs(latitude)+(1\|grouping)** | **5** | **49** | **645.2** |
| **2. elevation*abs(latitude)+(elevation\|grouping)** | **7** | **25** | **645.9** |
| 3. elevation*abs(latitude)+(elevation*abs(latitude)\|grouping) | 7 | 10 | 646.6 |
| 4. elevation*abs(latitude)*cloud_cover+(1\|grouping) | 9 | 19 | 647.6 |

**Table S6. Significant effect of elevation across latitude on pollination strategy** in the two best-fit models (Table S5) of the subset data; the last two columns indicate how many (across 100) random subsets yielded significant p-values.

| model | estimate | z-value | p-value | <0.05 | <0.01 | AUC |
|---|---|---|---|---|---|---|
| 1. elevation*abs(latitude) | -0.781 | -3.973 | 0.001 | 100 | 97 | 0.805 |
| 2. elevation*abs(latitude) | -0.715 | -3.339 | 0.016 | 94 | 82 | 0.814 |

**Table S7. Results from phylogenetic generalized linear mixed models on pollination mode** (assuming Brownian motion), showing a significant effect of elevation and latitude on pollination mode, with higher probability of vertebrate pollination at high elevations in the tropics than in the temperate zone. PGLMM simultaneously estimates the strength of phylogenetic signal in the residuals and performs a likelihood ratio test on the hypothesis that there is no phylogenetic signal in the data.

| | Value | Std-error | z-score | p-value |
|---|---|---|---|---|
| **full model**: *phylogenetic signal 10.62, p-value <0.001* | | | | |
| **(Intercept)** | -0.635 | 1.181 | -0.537 | 0.591 |
| **elevation** | **1.354** | **0.268** | **5.041** | **<0.001** |
| **abs(latitude)** | 0.050 | 0.256 | 0.195 | 0.846 |
| **cloud cover** | -0.015 | 0.248 | -0.059 | 0.953 |
| **elevation:abs(latitude)** | **-0.950** | **0.248** | **-3.835** | **<0.001** |
| **elevation: cloud cover** | -0.147 | 0.245 | -0.602 | 0.547 |
| **abs(latitude): cloud cover** | -0.051 | 0.229 | -0.223 | 0.824 |
| **elevation:abs(latitude): cloud cover** | 0.071 | 0.231 | 0.305 | 0.760 |
| **reduced model**: *phylogenetic signal: 10.02, p-value <0.001* | | | | |
| (Intercept) | -0.682 | 1.139 | -0.599 | 0.550 |
| **elevation** | **1.234** | **0.211** | **5.852** | **<0.001** |
| abs(latitude) | 0.114 | 0.224 | 0.508 | 0.611 |
| **elevation:abs(latitude)** | **-0.855** | **0.190** | **-4.497** | **<0.001** |

**Most important literature for pollinator data (Table S1)**


Abrahamczyk, S. & Renner, S.S. (2015). The temporal build-up of hummingbird/plant mutualisms in North America and temperate South America. *BMC Evol Biol,* 15, 104.

Abrahamczyk, S., Souto-Vilarós, D., Renner, S.S. (2014). Escape from extreme specialization: passionflowers bats and the sword-billed hummingbird. *Proc R Soc B*, 281, 20140888.

Ackermann, M. & Weigend, M. (2006). Nectar, Floral Morphology and Pollination Syndrome in Loasaceae subfam. Loasoideae (Cornales). *Annals of Botany,* 98, 503–514.

Aguilar-Rodríguez, P.A., Krömer, T., Tschapka, M., García-Franco, J.G., Escobedo-Sarti, J., MacSwiney, G.M.C. (2019). Bat pollination in Bromeliaceae. *Plant Ecology & Diversity*. https://doi.org/10.1080/17550874.2019.1566409

Alcantara, S. & Lohmann, L.G. (2010). Evolution of floral morphology and pollination system in Bignonieae (Bignoniaceae). *American Journal of Botany,* 97, 782-796.

Amico, G.C., Vidal-Russell, R., Nickrent, D.L. (2007). Phylogenetic relationships and ecological speciation in the mistletoe *Tristerix* (Loranthaceae): the influence of pollinators, dispersers, and hosts. *Am. J. Bot.,* 94, 558-567.

Castro, C.C. & Araujo, C.A. (2004). Distyly and sequential pollinators of *Psychotria nuda* (Rubiaceae) in the Atlantic rain forest, Brazil. *Plant Systematics and Evolution*, 244, 131–139.

Dellinger, A.S., Pérez-Barrales, R., Michelangeli, F.A., Penneys, D.S., Fernández-Fernández, D.M., Schönenberger, J. (2021). Low bee visitation rates explain pollinator shifts to vertebrates in tropical mountains. *New Phyt,* 231, 864-877.

Emms, S.K., Arnold, M.L. (2000). Site-to-site differences in pollinator visitation patterns in a Louisiana iris hybrid zone. *Oikos*, 91, 568-578.

Ferreira, C., Maruyama, P.K., Oliveira, P.E. (2016). Convergence beyond flower morphology? Reproductive biology of hummingbird-pollinated plants in the Brazilian Cerrado. *Plant Biol J*, 18, 316-324.

Givnish, T.J*.,* Barfuss, M.H*.,* Van Ee, B*.,* Riina, R*.,* Schulte, K*.,* Horres, R*., et al. (*2014)*.* Adaptive radiation, correlated and contingent evolution, and net species diversification in Bromeliaceae*. Mol. Phylogenet. Evol*., 71,55–78*.*

Grant, V., & Grant, K.A. (1965). Flower pollination in the phlox family. New York, Columbia University Press.

Grant, V. (1994). Modes and origins of mechanical and ethological isolation in angiosperms. *PNAS*, 91, 3-10.

Chuang, T.I., & Heckard, L.R. (1992). A taxonomic revision of *Orthocarpus* (Scrophoulariaceae – tribe Pedicularieae). Systematic Botany, 17, 560-582.

Karron, J., Thumser, N., Tucker, R., Hessenauer, A.J. (1995). The influence of population density on outcrossing rates in *Mimulus ringens*. *Heredity*, 75, 175–180.

Kriebel, R., Drew, B., González-Gallegos, J.G., Celep, F., Heeg, L., Mahdjoub, M.M., Sytsma, K.J. (2020). Pollinator shifts, contingent evolution, and evolutionary constraint drive floral disparity in *Salvia* (Lamiaceae): Evidence from morphometrics and phylogenetic comparative methods. *Evolution*, 74, 1335-1355.

Lagomarsino, L.P., Forrestel, E.J., Muchhala, N., Davis, C.C. (2017). Repeated evolution of vertebrate pollination syndromes in a recently diverged Andean plant clade. *Evolution*, 71, 1970-1985.

Mesquita-Neto, J.N., Silva-Neto, C.M., Franceschinelli, E.V. (2015). Theoretical predictions of plant-pollinator interactions in sympatric species of *Psychotria* (Rubiaceae) in Cerrado of Brazil. Plant *Ecology and Evolution*, 148, 229-236.

Meudt, H.M. & Simpson, B.B. (2007). Phylogenetic analyses of morphological characters in *Ourisia* (Plantaginaceae): taxonomic and evolutionary implications. Ann Miss Bot Gard, 94(3), 554-570.

Ocampo Pérez, J., Coppens d'Eeckenbrugge, G. (2017). Morphological characterization in the genus *Passiflora* L.: an approach to understanding its complex variability. *Plant Syst Evol* 303, 531–558.

Pérez, F., Arroyo, M.T., Medel, R., Hershkovitz, M.A. (2006). Ancestral reconstruction of flower morphology and pollination systems in *Schizanthus* (Solanaceae). *Am J Bot*, 93(7), 1029-38.

Rosas-Guerrero, V., Quesada, M., Armbruster, W.S., Pérez-Barrales, R. and Smith, S.D. (2011). Influence of pollination specialization and breeding system on floral integration and phenotypic variation in *Ipomoea*. *Evolution*, 65, 350-364.

Sakai, S., & Wright, S. (2008). Reproductive ecology of 21 coexisting *Psychotria* species (Rubiaceae): when is heterstyly lost? *Biol J Linn Soc*, 93, 125-134.

Serrano-Serrano, M.L., Rolland, J., Clark, J.L., Salamin, N., Perret, M. (2017). Hummingbird pollination and the diversification of angiosperms: an old and successful association in Gesneriaceae. *R Soc Proc B*, 284, 20162816.

Smith, S.W., Ané, C., Baum, D.A. (2008). The role of pollinator shifts in the floral diversification of Iochroma (Solanaceae). *Evolution*, 62, 793-806.

Tripp, E.A., & Manos, P.S. (2008). Is floral specialization an evolutionary dead-end? Pollination system transitions in *Ruellia* (Acantahceae). *Evolution,* 62, 1712-1737.

Tripp, E.A., & Tsai, Y.H.E. (2017). Disentangling geographical, biotic, and abiotic drivers of plant diversity in neotropical *Ruellia* (Acanthaceae). *PLoS ONE*, 12, e0176021.

Vargas, O.M., Goldston, B., Grossenbacher, D.L. and Kay, K.M. (2020), Patterns of speciation are similar across mountainous and lowland regions for a Neotropical plant radiation (Costaceae: *Costus*). *Evolution*, 74, 2644-2661.

von Hagen, B.K., & Kadereit, J.W. (2001). The phylogeny of *Gentianella* (Gentianaceae) and its colonization of the southern hemisphere as revealed by nuclear and chloroplast DNA sequence variation. *Organisms Diversity & Evolution*, 1, 61-79.

Wester, P., & Claßen-Bockoff, R. (2011). Pollination syndromes of new world *Salvia* species with special reference to bird pollination. *Ann Miss Bot Gard*, 98, 101-155.

Whittal, J.B., & Hodges, S.A. (2007). Pollinator shifts drive increasingly long nectar spurs in columbine flowers. *Nature*, 447, 706-709.

Wilson, P., Wolfe, A.D., Armbruster, W.S. and Thomson, J.D. (2007), Constrained lability in floral evolution: counting convergent origins of hummingbird pollination in *Penstemon* and *Keckiella*. *New Phytologist*, 176, 883-890.
